# FastCAR: Fast Correction for Ambient RNA to facilitate differential gene expression analysis in single-cell RNA-sequencing datasets

**DOI:** 10.1101/2022.07.19.500594

**Authors:** Marijn Berg, Ilya Petoukhov, Inge van den Ende, Kerstin B. Meyer, Victor Guryev, Judith M. Vonk, Orestes Carpaij, Martin Banchero, Rudi W. Hendriks, Maarten van den Berge, Martijn C. Nawijn

## Abstract

Cell type-specific differential gene expression analyses based on single-cell transcriptome datasets are sensitive to the presence of cell-free mRNA in the droplets containing single cells. This so-called ambient RNA contamination may differ between samples obtained from patients and healthy controls. Current ambient RNA correction methods were not developed specifically for single-cell differential gene expression (sc-DGE) analyses and might therefore not sufficiently correct for ambient RNA-derived signals. Here, we show that ambient RNA levels are highly sample-specific. We found that without ambient RNA correction, sc-DGE analyses erroneously identify transcripts originating from ambient RNA as cell type-specific disease-associated genes. We therefore developed a computationally lean and intuitive correction method, Fast Correction for Ambient RNA (FastCAR), optimized for sc-DGE analysis of scRNA-Seq datasets generated by droplet-based methods including the 10XGenomics Chromium platform. FastCAR uses the profile of transcripts observed in libraries that likely represent empty droplets to determine the level of ambient RNA in each individual sample, and then corrects for these ambient RNA gene expression values. FastCAR can be applied as part of the data pre-processing and QC in sc-DGE workflows comparing scRNA-Seq data in a health versus disease experimental design. We compared FastCAR with two methods previously developed to remove ambient RNA, SoupX and CellBender. All three methods identified additional genes in sc-DGE analyses that were not identified in the absence of ambient RNA correction. However, we show that FastCAR performs better at correcting gene expression values attributed to ambient RNA, resulting in a lower frequency of false-positive observations. Moreover, the use of FastCAR in a sc-DGE workflow increases the cell-type specificity of sc-DGE analyses across disease conditions.

## Introduction

Single cell RNA sequencing (scRNA-Seq) is revolutionizing basic and translational biomedical research. The ability to quantify RNA expression in individual cells with high throughput enables quantification of cell type composition of complex tissue samples and characterization of their transcriptional phenotypes, or cell states, in great detail^1^. The use of scRNA-Seq to compare healthy and diseased tissue samples can reveal differences in cell type proportions and identify unique, disease-associated cell types, cell states, cell-cell interactions or cell-state transitions, all of which can be used to chart the pathogenesis of disease at unprecedented level^2^. In such efforts, scRNA-seq can be applied to perform cell type-specific (or single-cell) differential gene expression (sc-DGE) analyses between healthy and diseased tissue samples.

We previously published a comparison of the cellular landscape in airway wall samples between healthy controls and patients with asthma^3^. While comparing cell-type composition is relatively straightforward, we observed that sc-DGE analyses between healthy and diseased tissue samples frequently yielded identification of differentially expressed genes in cell types that were unlikely to express these genes, indicating that the observed gene expression probably originated from the ambient RNA. This despite having corrected for ambient RNA using SoupX^4^.

Ambient RNA is cell-free mRNA that is released during preparation of single-cell suspensions for scRNA-Seq analysis and is one of the features that limits sc-DGE analyses, next to sparsity of data and the presence of doublets^5,6^. Sc-DGE analysis methods that take into account the sparsity of data or use pseudo-bulk approaches per cell type are being developed^7^. Doublets can be transcriptionally identified and removed from the dataset during pre-processing of scRNA-Seq data^6^. The ambient RNA present in the cell suspension will be captured by all beads during cell partitioning in droplet-based scRNA-Seq methods, irrespective of the presence or absence (‘empty’ droplets) of a cell. Consequently, cell-type specific mRNA released into the ambient RNA will also be detected at low levels in cell types that do not express this gene natively.

The composition of ambient RNA depends on the cell type composition and processing of the tissue, and is therefore highly sample-specific. When comparing gene expression profiles across cell types within a single sample, transcripts of ambient RNA will be shared and will not be identified as differentially expressed genes. In contrast, when comparing gene expression profiles in a cell-type specific fashion between different samples, the ambient RNA composition might be different between the samples. In such a case, transcripts identified as differentially expressed genes may be derived from both cellular RNA and contaminating ambient RNA, leading to false-positive results. Considering our previous results with SoupX a new method for ambient RNA correction that allows sc-DGE studies comparing healthy to diseased tissue samples is urgently needed.

Here, we characterized the contamination by ambient RNA in sc-DGE analyses, and present a novel method ‘FastCAR’ (Fast Correction for Ambient RNA) to quickly identify and correct for ambient RNA in droplet-based scRNA-Seq data. We provide a rationale for selection of the genes that should be corrected on the basis of the data retained within the gene expression matrix, without the need for prior knowledge on the expected cell-type specific gene expression patterns. Furthermore we compare its performance to the other ambient RNA correction methods SoupX and CellBender-remove-background which were either not thorough enough or computationally prohibitive^8^. The use of FastCAR as part of the scRNA-seq data pre-processing workflow allows for more accurate sc-DGE analyses between disease conditions or other experimental groups.

## Materials and Methods

### scRNA-seq datasets

#### Bronchial biopsies from healthy controls and asthma patients

To test FastCAR, we used our previously published^3^ scRNA-Seq dataset obtained from bronchial biopsies of six asthma patients and six healthy controls. Here, we used the same cells as in our previous study^3^, however using an updated cell-type annotation to better reflect our current understanding of the data. Mapping and counting was performed using 10x Genomics Cell Ranger 3.1.0 with the GRCh38 genome reference and gene annotation from Ensembl release 93 to generate new count matrices for these barcodes, and without the ambient RNA correction using SoupX that was applied in the original dataset^3^.

#### PBMCs from healthy donors and COVID-19 patients

As another disease/control dataset we used PBMCs from seven healthy donors and seven hospital admitted COVID-19 patients.^9^ These were processed on the SeqWell^10^ platform.

#### Differential gene expression analyses

To perform the differential expression analyses we used R package EdgeR^11^ using the likelihood ratio test on pseudo-bulk per cell type, using the disease condition as the contrast between groups. The aggregate pseudo-bulk count matrices were generated per sample and cell type using the ‘PseudobulkExpression’ function of Seurat 4.02^12^.

#### Ambient RNA removal by ‘SoupX’^4^

We applied SoupX using the suggested settings by the *autoest* function of the original tutorial starting from the count matrices. The cell selection step was not applied as we used only the libraries that were selected as live, high-quality cells in our previous publication^3^. For the cluster annotations we used the updated cell labels reflecting our current understanding of gene expression in different cell types.

#### Ambient RNA removal by ‘CellBender remove background’^8^

To perform the ambient RNA correction using CellBender remove background we used the default recommended settings (fpr 0.01, epochs 150). The cell selection was not applied as we used only the libraries that were selected as live, high-quality cells in our previous publication^3^.

#### Seurat

Processing of the data was done using the R package Seurat version 4.02^12^

#### R

Data processing was performed using R version 4.1.2

## Results

In order to develop a method that can be used to correct for ambient RNA in a sc-DGE analysis, we took advantage of a previously reported data set in which we reported the changes in the cellular landscape of the airway wall in patients with asthma compared to healthy controls^3^. After updating the cell labels to better reflect current understanding (Fig 1A) we found that transcripts of several cell type-specific genes were present in other cell types, not known to express these genes, albeit at much lower levels than the actual expressing cell type (Fig 1B). These included for example, *SCGB3A1*, expressed in secretory cells^13,14^, *IGKC* from B cells^15,16^ and *HBB*, originating from erythrocytes^16,17^ which while not identified as distinct cell types in the data would have been present in the biopsy. Moreover, these genes were all identified as being significantly differentially expressed between asthma patients and healthy controls. We hypothesized that such ‘ectopic’ DE gene expression patterns could be attributed to differences in ambient RNA that were not fully corrected for.

**Figure 1:**
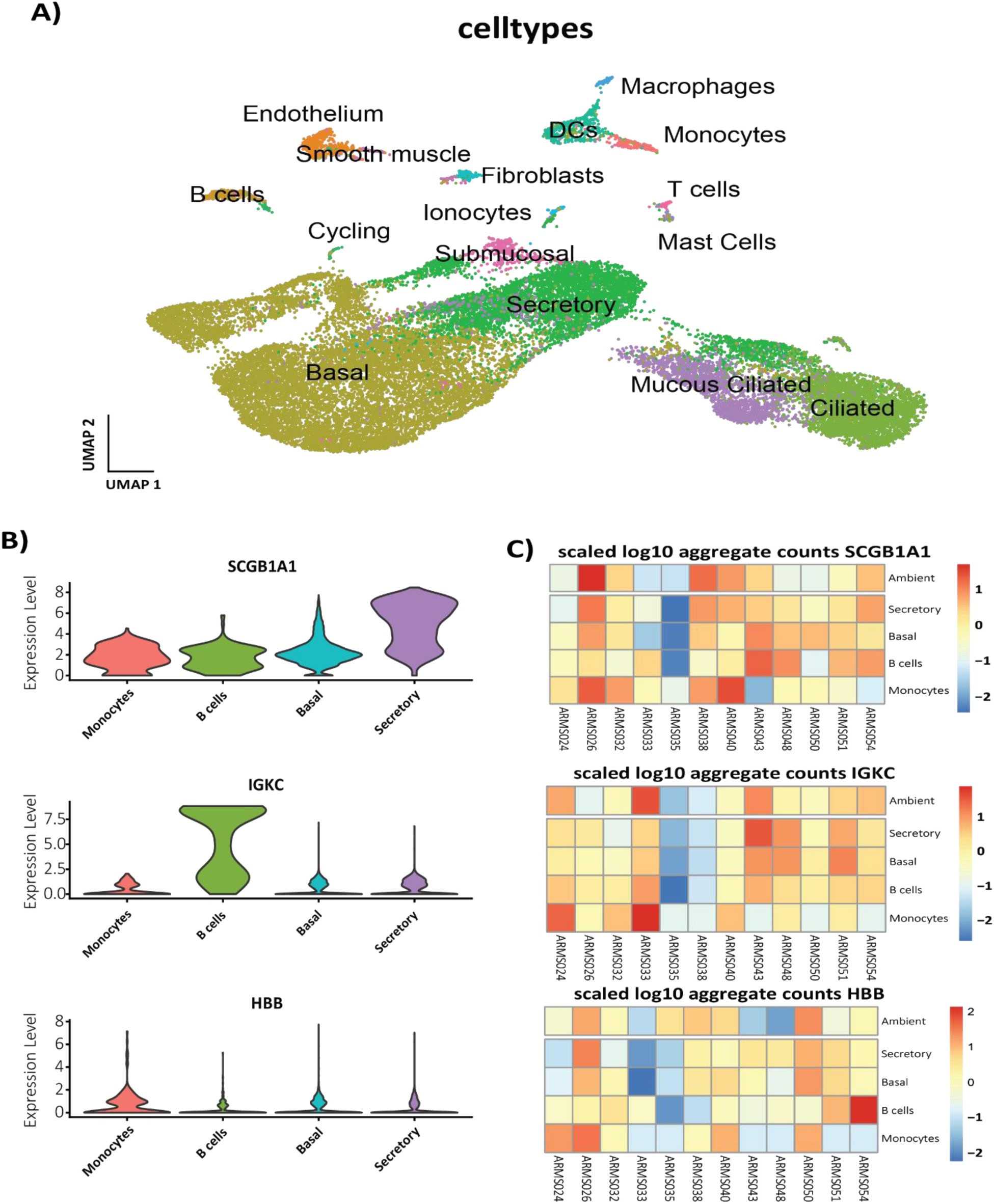
Ambient RNA is sample specific: **A)** UMAP of the associated data **B)** Violin plots of the mRNA levels of selected cell type markers found in the ambient RNA in selected cell types. **C)** Heatmaps of the scaled log10 aggregate pseudo bulk expression of selected genes in samples from the bronchial biopsy dataset in selected cell types and the ambient [nUMI <=100] libraries.

This ambient RNA is sample-specific which leads to samples where the total ambient RNA (summed UMI in libraries with =< 100 UMI) of a gene is higher to also have relatively higher presence of that gene in non-expressing cells compared to other samples (Fig 1C). This sample-specific profile of gene ‘expression’ is what is used to correct for this ambient RNA and make sc-DGE more accurate.

### FastCAR algorithm

FastCAR determines the ambient RNA profile to correct the cell expression for on a gene by gene basis. The user provides a threshold for the number of Unique Molecular Identifiers (UMI) per sequencing library (***thE***), every library (***j***) with that number of UMIs or fewer is used to generate the ambient RNA profile. For every gene (***g***), the fraction of these selected libraries containing any UMIs of that gene (***frC***) is determined, as well as the highest number of UMIs of that gene occurring in a single library (***gMax***). If ***frC*** exceeds the user provided allowable fraction of ambient affected cells (***frAA***), the UMI counts for that gene in each cell is reduced by ***gMax***. If this results in negative counts, the number of counts of that gene in the cell is set to 0.

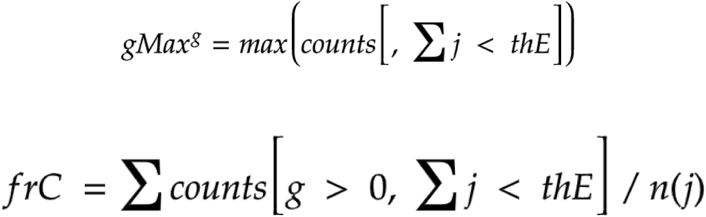

DGE methods for scRNA-seq data use a cut-off for the minimum number of cells that need to be expressing a gene in a sample and cluster before it is considered for testing, ***frAA*** can be set based on this by choosing a fraction that matches this cut-off. ***thE*** can be set by default to 100 UMI but more informed choices lead to better results as explained in the workflow example shown in figure 2 and explained in the results.

**Figure 2:**
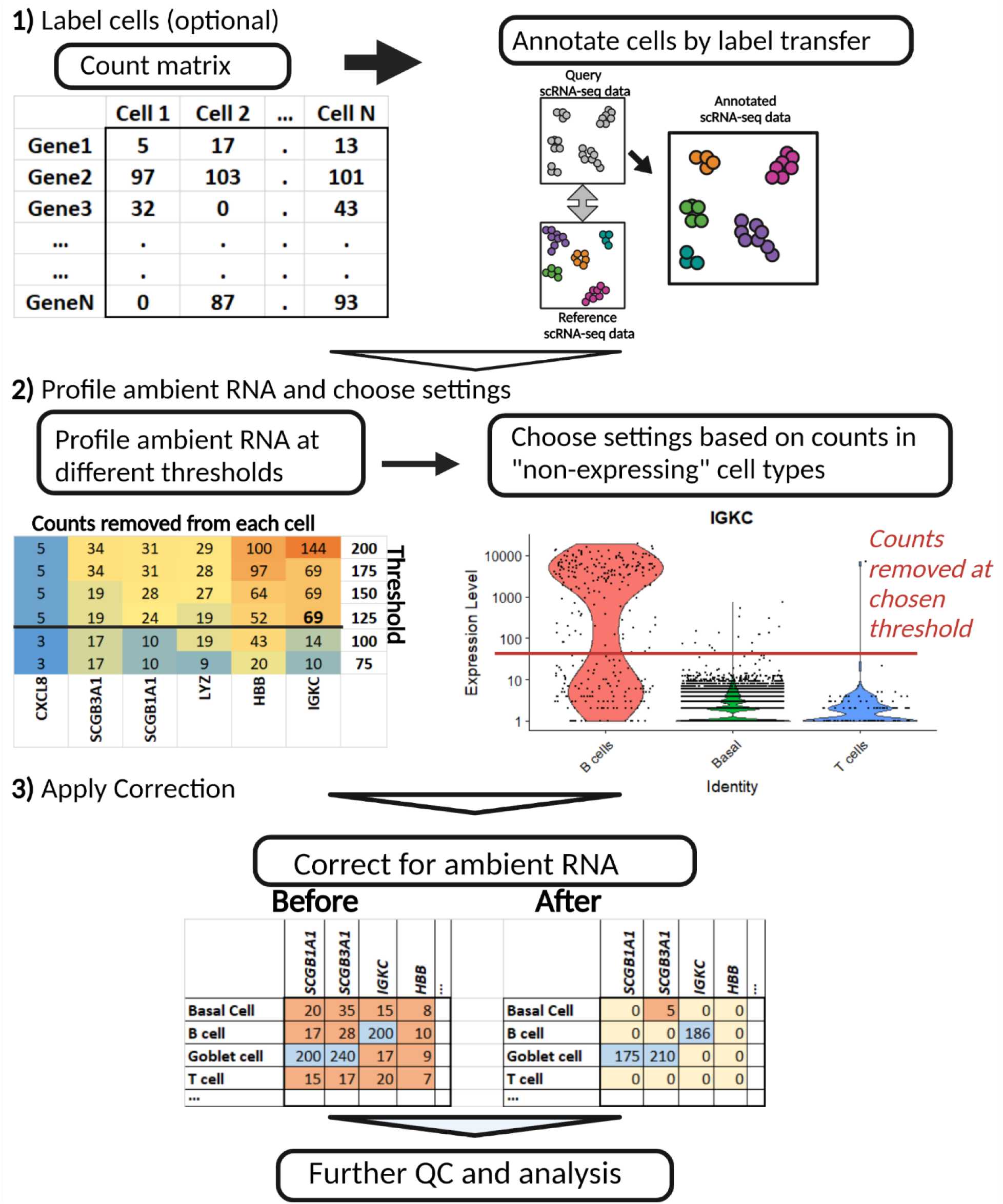
Workflow to select appropriate settings and apply FastCAR to a scRNA-seq dataset.

### FastCAR: methodology and setting thresholds

We developed a method named FastCAR for optimized correction of ambient RNA levels to allow more effective sc-DGE analysis in studies comparing healthy and diseased samples and similar experimental designs. FastCAR uses the absolute number of UMIs from the ambient-RNA containing libraries to profile the gene expression pattern and levels present in ambient RNA and perform a correction for these levels in all libraries that are identified as live, single cells.

As stated earlier, there are two variables to consider when running FastCAR. Firstly, the minimum allowable fraction of libraries that contains ambient RNA of each gene. (***frAA***). Secondly, the maximum UMI-per-library threshold at which libraries are considered to only contain ambient RNA(***thE***).

The minimum allowable fraction of cells affected by the ambient RNA that needs to be corrected for depends on the method of sc-DGE analysis that will be performed. Most sc-DGE methods apply a threshold for the minimum fraction of cells per cluster that need to express a gene to accept it as a DE gene. Therefore, all genes that are found to be present in a lower proportion of the ambient RNA libraries (at the chosen empty library threshold) will not be identified in the sc-DGE method, and therefore do not need to be corrected for. The default setting for this contamination chance parameter is 0.005; any genes present in less than 5 out of each 1000 ambient RNA libraries are ignored. This limits the total profile of ambient RNA that needs to be corrected for quite significantly.

The threshold for what is considered an empty library is often arbitrarily set at 100 UMIs/library^8^. We have established a workflow (Figure 2) that illustrates a method to facilitate choosing an appropriate threshold using known cell labels and expected cell type-specific genes. If a reference is available for the tissue then transfer learning with for instance scArches^18^ can provide these labels. Methods to assist in setting these thresholds without cell labels are described on the GitHub page.

The next step is to profile the level of RNA that will be removed if the threshold for an empty library is increased. The higher the threshold, the higher the number of counts that are removed of each gene as can be seen in the heatmap. Whether the correction level of genes is adequate can be gauged by comparing the number of counts that will be removed at a threshold, to the level of expression in cells that are expected to express them, and the levels at which it is present in non-expressing cells as shown in the violin plot. A proper threshold will help to mostly remove transcripts of genes likely to affect DGE analysis while otherwise removing as little transcripts/signals as possible. As such the recommended threshold is one where the number of removed counts per cell is around the actual level observed in non-expressing cells. In the figure *IGKC* is used as an example to choose a threshold. At an empty library threshold of 125 UMIs, the counts in the non-expressing libraries will be mostly corrected for while minimally affected the expressing cells.

In general, a higher threshold will result in more genes being selected for correction and a more extensive removal of ambient RNA-derived transcripts. However such a threshold will also lower the expression values of commonly expressed genes including the mitochondrial genes and could overcorrect genes that are highly expressed in some cell types but lowly expressed in others, completely removing the measured expression in the latter. If the threshold is set to levels over about 500, lowly expressing but live cells such as T cells may be included in the “ambient” profile, this could raise ***gMax*** for those genes and overcorrect the expression such that none will remain in these cell types.

After the threshold has been set FastCAR can be applied to all samples and the resulting count matrices can be used in downstream analyses.

### Effect of FastCAR correction on sc-DGE results

We profiled the ambient RNA of our bronchial biopsy dataset using a threshold of 150 UMIs/library for ambient RNA and minimal allowable affected fraction of 0.005. Many of the genes identified to be part of the ambient RNA using these settings were found to be differentially expressed in more than two cell types between asthma and control. After applying FastCAR and correcting the expression of these genes, we found that many of the previously significant DGE results were no longer observed (Fig 3A).

**Figure 3:**
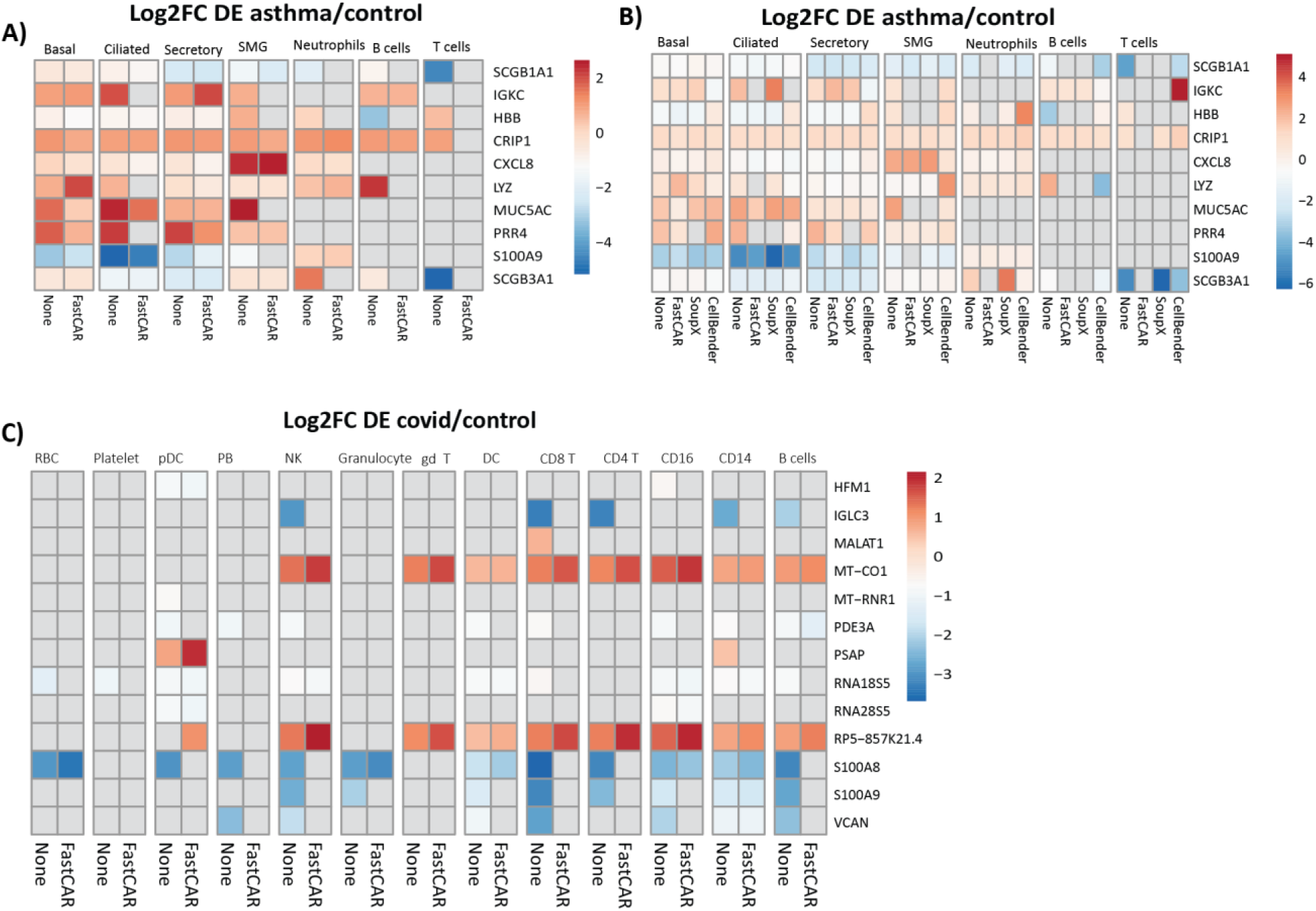
Effect of applying FastCAR and comparison to other methods. **A)** Log2 fold change of selected significant DE genes between asthma and control in bronchial biopsies before and after being corrected for by FastCAR. **B)** Log2 fold change of selected significant DE genes between asthma and control in bronchial biopsies before and after being corrected for by FastCAR, SoupX and Cellbender. **C)** Log2 fold change of selected significant DE before and after being corrected for by FastCAR for genes that were corrected for in at least one sample and found to be differentially expressed in PBMCs between COVID-19 patients and healthy controls.

In total, across all cell types, 372 out of 5067 identified DGE lost significance in the sc-DGE analysis in one or more cell types after FastCAR correction (Table 2). Next to that, 214 genes were identified as differentially expressed only after correction.

### Comparison of FastCAR to other methods

Next, we compared FastCAR to other available ambient RNA correction methods, CellBender^8^ and SoupX^4^ (Fig 3B).

When comparing the total number of significant DGE results across all cell types of genes that were corrected for by FastCAR in at least one of the samples we observe that these are most reduced by FastCAR (table 1).

**Table 1:** Number of times that genes corrected for by FastCAR in at least one sample were significantly differentially expressed in cell types before correction and after applying different ambient RNA corrections.

|  | uncorrected | FastCAR | CellBender | SoupX |
| --- | --- | --- | --- | --- |
| **Secretory** | 49 | 47 | 47 | 49 |
| **Submucosal** | 20 | 14 | 45 | 18 |
| **Ciliated** | 40 | 46 | 39 | 34 |
| **Basal** | 36 | 38 | 42 | 39 |
| **Mucous Ciliated** | 161 | 154 | 167 | 161 |
| **T cells** | 21 | 13 | 7 | 18 |
| **Endothelium** | 29 | 21 | 22 | 20 |
| **Fibroblasts** | 13 | 6 | 10 | 12 |
| **DCs** | 16 | 11 | 15 | 12 |
| **Ionocytes** | 12 | 21 | 28 | 13 |
| **B cells** | 19 | 9 | 27 | 15 |
| **Neutrophils** | 15 | 17 | 11 | 13 |
| **Macrophages** | 20 | 23 | 13 | 24 |
| **Cycling** | 23 | 15 | 13 | 22 |
| **Smooth muscle** | 5 | 4 | 4 | 4 |
| **Mast Cells** | 19 | 13 | 9 | 19 |
| **Total** | **498** | **452** | **499** | **473** |

There are also differences that FastCAR may not correct for, therefore we compared the total number of significant sc-DE genes and the effects on the different ambient RNA correction methods (table 2). We found large differences between the results before and after the various corrections. CellBender showed the least overlap between the corrected and uncorrected datasets but also large number of new DEGs and one DEG which effect is inverted after correction.

**Table 2:**
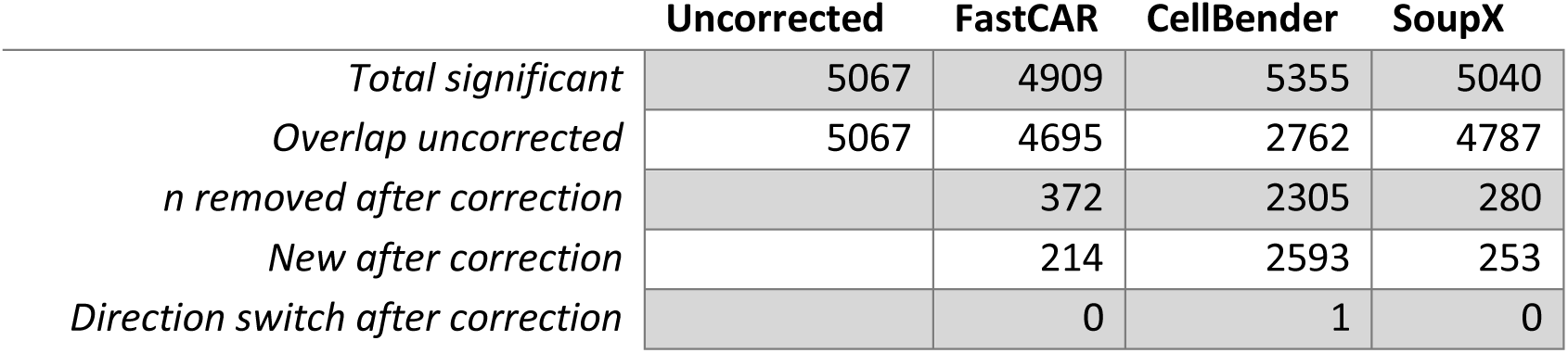
total number of sc-DGE results with different corrections and the comparison to the uncorrected results

|  | Uncorrected | FastCAR | CellBender | SoupX |
| --- | --- | --- | --- | --- |
| *Total significant* | 5067 | 4909 | 5355 | 5040 |
| *Overlap uncorrected* | 5067 | 4695 | 2762 | 4787 |
| *n removed after correction* |  | 372 | 2305 | 280 |
| *New after correction* |  | 214 | 2593 | 253 |
| *Direction switch after correction* |  | 0 | 1 | 0 |

To determine the difference in correction between methods for individual genes we plotted the expression of *IGKC* and *SCGB1A1* without correction and after applying the different corrections. method (supplementary figure 2 A/B) For both of these genes the ambient RNA is more completely removed by FastCAR than the other methods.

#### Comparison to other methods

We next asked whether correcting for ambient RNA by FastCAR could negatively affect downstream processing of the data, such as clustering. To this end, we compared the clusters identified in our bronchial biopsy dataset by Seurat both before and after ambient RNA correction. Identical settings and cells were used in both clustering efforts, normalization and scaling was performed using SCTransform while regressing out the percentage mitochondrial RNA, and the first ten principal components were used for the clustering at resolution 0.1. The libraries with fewer than 150 UMIs were used to profile the ambient RNA. Gene expression levels were corrected for maximal expression in the empty droplets if they had a minimal allowable fraction size higher than 0.005. We used the Jaccard index to compare whether the cells cluster similarly before and after ambient RNA correction and found that clustering and cell type identification is not strongly affected by the presence of ambient RNA as shown in supplementary figure 1. The main changes that occur after ambient RNA correction are observed in clusters identified along a differentiation trajectory lacking discrete cell type transitions, such as the basal and secretory cells of the airway epithelium where the cells are relatively arbitrarily split into separate clusters.

To validate whether FastCAR also works on other datasets and platforms we used the CoVID-19/healthy control PBMC dataset from Wilk et al^9^(Fig 3C), here too we observed differentially expressed genes that might result from ambient RNA have smaller effect sizes and are often no longer significant after correction with FastCAR.

FastCAR is an effective method to correct for ambient RNA that is most likely to affect DGE analyses between groups within cell types in scRNA-seq studies, applicable to multiple droplet-based methods and compatible with downstream analyses.

## Discussion

We have developed FastCAR (Fast Correction for Ambient RNA), a computational method that provides an unbiased way to identify and correct for the ambient RNA likely to affect DGE analyses on a per-sample basis that is intuitive and easy-to-use. FastCAR identifies the genes in the ambient RNA and applies a threshold for filtering that can be adapted to the settings of the DGE analyses, effectively removing the ambient RNA from the DE gene results. We show that this method effectively removes ambient RNA, but still retains a large proportion (∼92%) of the DE genes observed prior to ambient RNA removal.

When comparing the results of DGE analysis before and after ambient RNA correction using FastCAR to the other methods we tested, there is a striking difference in the number of genes affected by correction procedures. Because FastCAR is made to only adjust for the expression genes likely to affect DGE analyses, it corrects only a small subset of genes and most results are identical to the uncorrected results. Both SoupX and CellBender adjust for the expression of many genes which results in large differences in the number and identity of the differentially expressed genes before and after ambient RNA correction. While both CellBender and FastCAR apply linear transformation of the data during correction, SoupX also applies a normalization that might interfere with certain downstream analyses. In the absence of a gold standard to compare the results of the DGE analyses and the low sample numbers in the dataset we used for testing, it is not possible now to determine with certainty which of these methods is optimal for the identification of DE genes that reflect the biological truth. However, using well-established cell type-specific genes like *IGKC* and *SCGB1A1*, we could show that the performance for ambient RNA removal of these genes with FastCAR is superior compared to CellBender and SoupX. Other genes that FastCAR identifies and corrects for are also more thoroughly corrected for meaning that applying the other methods would still result on falsely identifying genes as differentially expressed that result from ambient RNA.

### Impact and possible uses

DGE analyses between healthy and control or other experimental groups in specific sub-populations of cells is a promising use of single cell data that may have large impact on our understanding of how the in situ behaviour of cells of the same type differs between groups. Correcting for the presence of ambient RNA will be vital to finding meaningful results in these analyses and FastCAR is an effective method to do so.

### Limitations

Because of the large variability in cells and processing there is no perfect threshold to use to profile the ambient RNA, or a method to determine with certainty what this threshold should be for a specific sample. This necessitates the use of arbitrary thresholds or user-defined thresholds. The FastCAR R package includes functions that help the user set these thresholds and allows for profiling the ambient RNA without performing the correction to facilitate choosing these thresholds.

A reasonable concern is whether removing the highest expression found in the ambient RNA is not too strict. We analysed the cell-type specific gene expression compared to the ambient expression levels and found these to be an order of magnitude higher. Consequently, the cell-type specific expression levels are well retained even for the genes that are highly expressed in the ambient RNA. Moreover, the FastCAR correction does not strongly affect clustering of the cells. This is not unexpected as the ambient RNA is a low and ubiquitous signal that has an equal chance of affecting each cell in a sample.

The correction method works under the assumption that cell containing libraries are equally likely to contain ambient RNA and that mRNA from lysed cells is the only meaningful source of transcripts that are not expressed in the measured cell. Other possible sources of such transcripts are barcode switching, where spontaneous errors in the cell barcodes cause transcripts to be assigned to the wrong cell. This may also be partially responsible for some of the signal^8^. FastCAR does not take such possibilities into account as it aims to remove just the genes most likely to affect DGE analyses, which is unlikely to occur as barcode switching is equally likely to affect each barcode.

In conclusion, to perform cell type-specific DGE analyses between groups there needs to be some correction to account for the sample-specific ambient RNA. FastCAR is a resource efficient method of thoroughly correcting the expression of genes most likely to be affected by ambient RNA. While other tools are available for general ambient RNA correction, FastCAR is more thorough in its correction than the other tested methods for genes it identifies as likely to affect sc-DGE.

## Code Availability

Instructions for installation and use of the FastCAR package as well as the source code are available online at https://github.com/Nawijn-Group-Bioinformatics/FastCAR

## Acknowledgements

M.C.N. acknowledges funding from the Netherlands Lung Foundation project no. 4.1.18.226

---

**Supplementary Figure 1:**
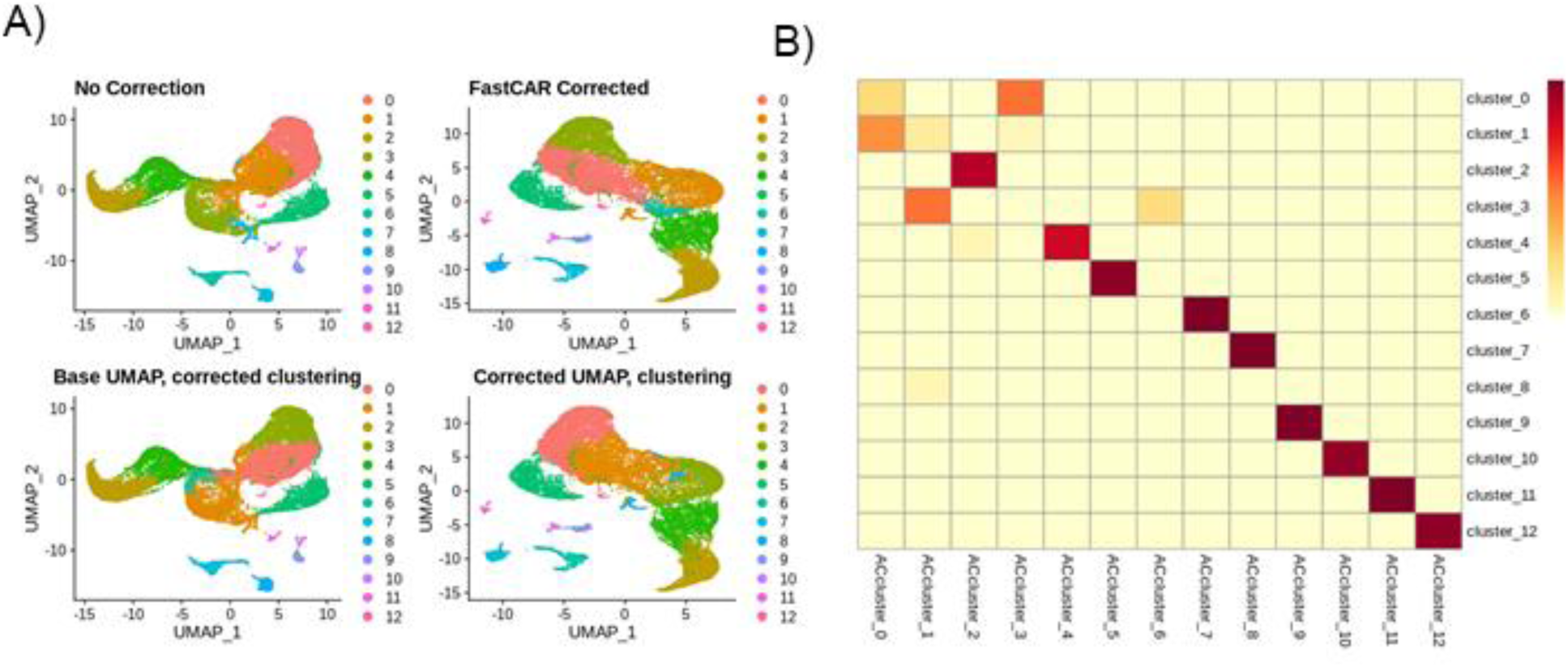
FastCAR correction does not strongly affect clustering. **A)** Effect of FastCAR on the UMAP and clustering in the same cells from bronchial biopsies. **B)** Jaccard index of the overlap of the cell contained in the clusters between corrected and non-corrected bronchial biopsies. The same cells cluster together even if the clusters get split differently along a gradient.

**Supplementary Figure 2:**
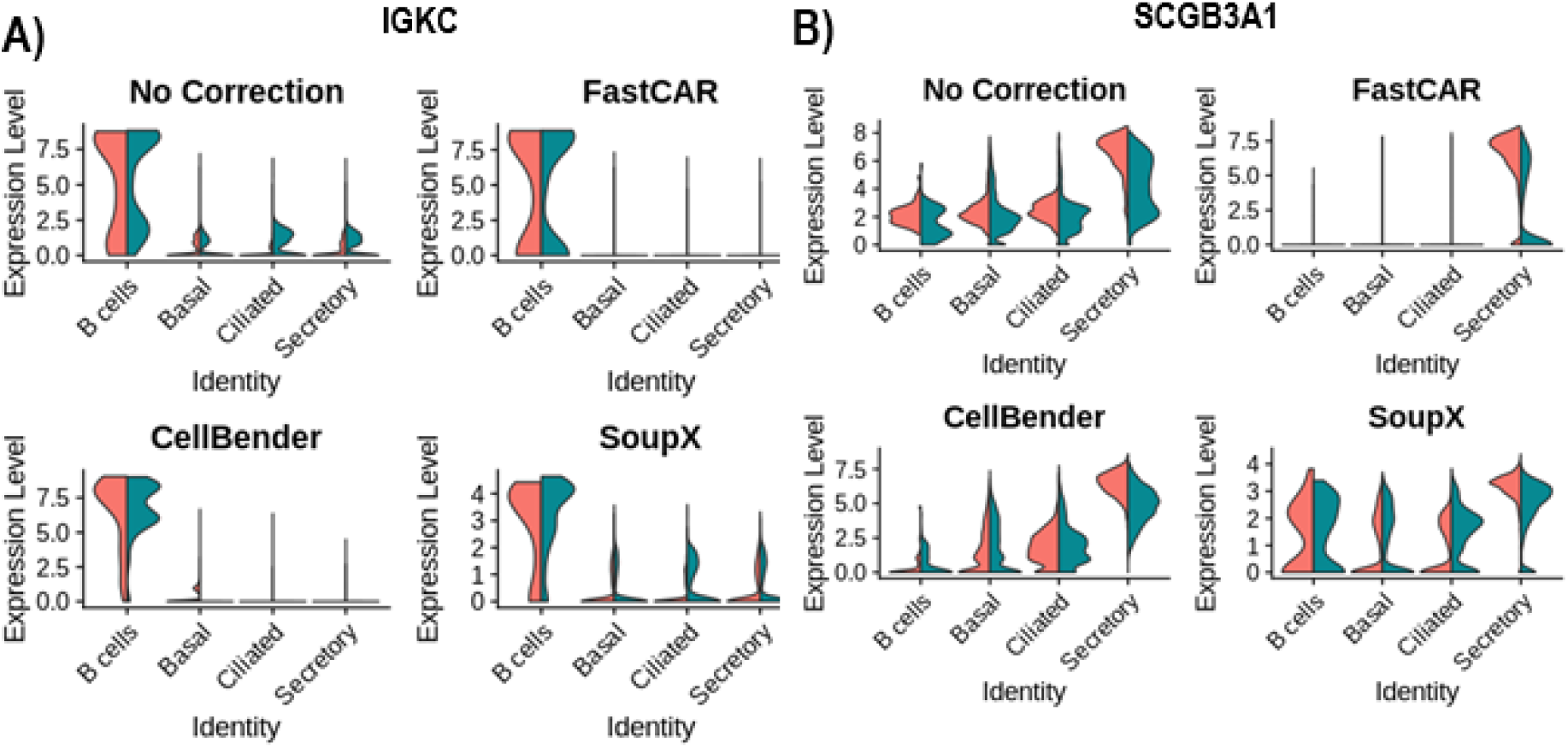
Comparison of applying different ambient RNA correction methods between asthma and control in selected cell types in bronchial biopsies. **A)** IGKC levels in selected expressing and non-expressing cell types without correction and after applying other correction methods. **B)** SCGB3A1 levels in selected expressing and non-expressing cell types without correction and after applying other correction methods.

## References

1. Macosko, E. Z. et al. Highly Parallel Genome-wide Expression Profiling of Individual Cells Using Nanoliter Droplets. Cell 161, 1202–1214 (2015).

2. Regev, A. et al. The Human Cell Atlas. eLife 6,.

3. Vieira Braga, F. A. et al. A cellular census of human lungs identifies novel cell states in health and in asthma. Nat. Med. 25, 1153–1163 (2019).

4. Young, M. D. & Behjati, S. SoupX removes ambient RNA contamination from droplet-based single-cell RNA sequencing data. GigaScience 9, giaa151 (2020).

5. Pierson, E. & Yau, C. ZIFA: Dimensionality reduction for zero-inflated single-cell gene expression analysis. Genome Biol. 16, 241 (2015).

6. Wolock, S. L., Lopez, R. & Klein, A. M. Scrublet: Computational Identification of Cell Doublets in Single-Cell Transcriptomic Data. Cell Syst. 8, 281-291.e9 (2019).

7. Wang, T., Li, B., Nelson, C. E. & Nabavi, S. Comparative analysis of differential gene expression analysis tools for single-cell RNA sequencing data. BMC Bioinformatics 20, 40 (2019).

8. Fleming, S. J., Marioni, J. C. & Babadi, M. CellBender remove-background: a deep generative model for unsupervised removal of background noise from scRNA-seq datasets. bioRxiv 791699 (2019) doi:10.1101/791699.

9. Wilk, A. J. et al. A single-cell atlas of the peripheral immune response in patients with severe COVID-19. Nat. Med. 26, 1070–1076 (2020).

10. Gierahn, T. M. et al. Seq-Well: portable, low-cost RNA sequencing of single cells at high throughput. Nat. Methods 14, 395–398 (2017).

11. Robinson, M. D., McCarthy, D. J. & Smyth, G. K. edgeR: a Bioconductor package for differential expression analysis of digital gene expression data. Bioinformatics 26, 139–140 (2010).

12. Hao, Y. et al. Integrated analysis of multimodal single-cell data. Cell 184, 3573-3587.e29 (2021).

13. Reynolds, S. D., Reynolds, P. R., Pryhuber, G. S., Finder, J. D. & Stripp, B. R. Secretoglobins SCGB3A1 and SCGB3A2 define secretory cell subsets in mouse and human airways. Am. J. Respir. Crit. Care Med. 166, 1498–1509 (2002).

14. Jackson, B. C. et al. Update of the human secretoglobin (SCGB) gene superfamily and an example of ‘evolutionary bloom’ of androgen-binding protein genes within the mouse Scgb gene superfamily. Hum. Genomics 5, 691 (2011).

15. Expression clustering - The Human Protein Atlas. https://www.proteinatlas.org/humanproteome/tissue/expression+cluster#cluster87.

16. Karlsson, M. et al. A single–cell type transcriptomics map of human tissues. Sci. Adv. 7, eabh2169.

17. Single cell type - HBB - The Human Protein Atlas. https://www.proteinatlas.org/ENSG00000244734-HBB/single+cell+type.

18. Lotfollahi, M. et al. Mapping single-cell data to reference atlases by transfer learning. Nat. Biotechnol. 1–10 (2021) doi:10.1038/s41587-021-01001-7.

